# Selective depletion of cancer cells with extrachromosomal DNA via lentiviral infection

**DOI:** 10.1101/2025.02.22.639631

**Authors:** Eunhee Yi, Amit D. Gujar, Dacheng Zhao, Kentaro Suina, Xue Jin, Katharina Pardon, Qinghao Yu, Larisa Kagermazova, Jef D. Boeke, Anton G. Henssen, Roel G.W. Verhaak

## Abstract

Dear Editor

Extrachromosomal DNA (ecDNA), a major focal oncogene amplification mode found across cancer, has recently regained attention as an emerging cancer hallmark^1,2^, with a pervasive presence across cancers^3,4^. With technical advancements such as high-coverage sequencing and live-cell genome imaging, we can now investigate ecDNA’s behaviors and functions ^5-7^. However, we still lack an understanding of how to eliminate ecDNA. We observed depletion of cells containing ecDNA during lentiviral but not transposon-based transduction while we sought to investigate the mechanism of ecDNA behavior. This discovery may provide critical information on utilizing a lentiviral system in emerging ecDNA research. Additionally, this observation suggests specific sensitivities for cells with ecDNA.

## Depletion of ecDNA-positive (ecDNA+) cell population after lentiviral transduction and selection for drug resistance

This phenomenon was first observed in PC3, a prostate cancer cell line model known to have extrachromosomal *MYC* amplification^8^. We sought to integrate multiple DNA constructs into the genomes of PC3 cells using a lentiviral transduction system. After three cycles of lentiviral transduction and antibiotic treatment, we observed a depletion of cells containing *MYC* ecDNAs, whereas cells containing linear *MYC* amplification, referred to as homogenously staining regions (HSRs), appeared unaffected (**Figure 1A and B**). We generated a methotrexate-resistant HeLa cell line (HeLa-MTX-Res) that develops dihydrofolate reductase (DHFR) ecDNAs. After the first lentiviral transduction with Cas9-expressing plasmid, we observed a significant increase in cells containing *DHFR* HSRs (75.8%) (**Supplementary Fig. 1A and B**). Two additional cycles of lentiviral transduction and antibiotic selection led to a complete depletion of cells containing *DHFR* ecDNAs (**Supplementary Fig. 1A and B**). Next, we checked whether antibiotic selection, carried out after lentiviral transduction, impacts the depletion of ecDNA+ cells. We found that puromycin treatment following lentiviral transduction had no further effect on the reduction of ecDNA+ cell populations or expansion of HSR+ cell populations in both PC3 (**Figure 1C**) and HeLa-MTX-Res cell line models (**Supplementary Fig. 1C**). Lentiviral transduction of smaller plasmids (<7kb) showed no effect on the depletion of the ecDNA+ populations, suggesting that lentiviral delivery of small molecules such as sgRNAs and shRNAs can be achieved (**Figure 1D**).

**Figure 1.**
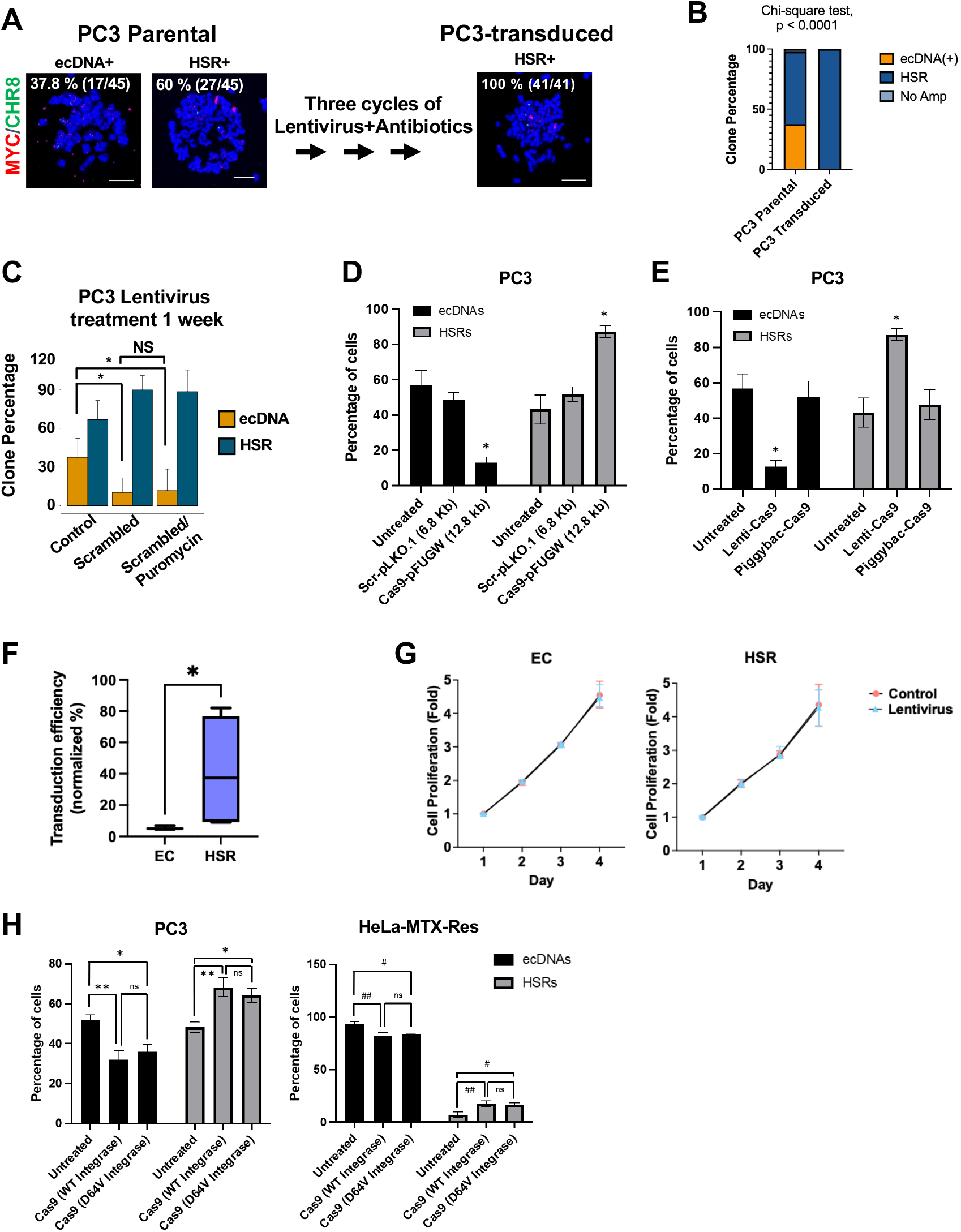
Lentiviral transduction causes the depletion of cancer cells with extrachromosomally amplified oncogenes. **A**. Oncogene amplification status before and after lentiviral transduction. Parental PC3 cells and the transduced PC3 cells that underwent three cycles of lentiviral transduction followed by antibiotics selection were synchronized at metaphase and processed for FISH analysis. MYC probe (red) and Chromosomal control probe (green) were used. The number of cells containing MYC amplification was quantified (n = 41). Bar = 10 micrometer. **B**. Graphical summary and statistical test of FISH analysis (Chi-square test, p < 0.0001). **C**. Switch of the subpopulation proportion after a single cycle of lentiviral transduction followed by puromycin treatment. PC3 cells were lentivirally transduced and selected with puromycin for 1 week. FISH analysis with MYC probe was performed to quantify subpopulations (T-test, *p < 0.05, n=3). **D**. Higher transgene/plasmid size in lentiviral construct depletes ecDNA+ cells with expansion of HSR+ cells. PC3 cells were either untreated or lentivirally transduced with indicated plasmids and selected with blasticidin for 2 weeks. FISH analysis with MYC probe was performed to quantify subpopulations (ANOVA, *p < 0.0001, n=3). **E**. Non-lentiviral (PiggyBac) plasmid does not deplete ecDNA+ cells. PC3 cells were either untreated or lentivirally transduced with Cas9 plasmid (Lenti-Cas9) or transfected with PiggyBac-Cas9 plasmid and selected with blasticidin for 2 weeks. FISH analysis with MYC probe was performed to quantify subpopulations (ANOVA, *p < 0.0001, n=3). **F**. Differential lentiviral transduction efficiency in PC3 single-cell clones with MYC-ecDNA (EC) or MYC-HSR (HSR). Two single-cell clones per each MYC amplification category were tested. Cells were transduced with lentivirus carrying green fluorescence protein (GFP) for 24 hours and subjected to flow cytometry analysis. Lentiviral transduction efficiency was calculated by quantifying the proportion of cells expressing GFP (T-test, *p < 0.05, n = 3). **G**. Cell proliferation comparison between PC3 single-cell clones with MYC-ecDNA (EC) or MYC-HSR (HSR). Two single-cell clones per each MYC amplification category were tested. Five sets of cells were transduced with lentivirus, and each set of cells was subjected to CellTiter-Glo assay on different days (posttransduction 1, 2, 3, 4, 5 days). The posttransduction proliferation rate was compared to the proliferation rate of the untreated control cells. **H**. Integrase-deficient lentivirus does not rescue the effect of integrase-intact lentivirus on ecDNA depletion. PC3-NCI and HeLa-MTX-Res cells were either untreated or transduced with Cas9-expressing lentivirus generated either using wild-type integrase (Cas9 WT Integrase) or point mutant integrase (Cas9 D64V Integrase) that does not integrate into the genome. Cells were cultured for 2 weeks. FISH analysis with oncogene probes was performed to quantify subpopulations (ANOVA *P < 0.01, **P < 0.05, #P < 0.05, ##P < 0.01, n = 2).

The PiggyBac transposon system, which integrates transgenes into host genomes via transposases, did not affect the depletion of ecDNA+ cell populations (**Figure 1E**). These results suggest that the lentiviral transduction method should be used with caution in ecDNA+ cells, especially those containing mixed subpopulations with different forms of oncogene amplification, as it alters the original ratio of cell subpopulations. Additionally, these results suggest the transposon system as an alternative way to deliver transgene to ecDNA+ cells without perturbing the ratio of cell subpopulations.

## Possible causes of lentiviral transduction mediated ecDNA+ cell depletion

Next, we sought to explain the relative depletion of ecDNA+ cells following lentiviral transduction. Before doing so, we ruled out the potential of massive parallel ecDNA reintegration into chromosomes after lentiviral transduction. The HSR pattern observed in the transduced cancer cells was almost identical to the preexisting HSR subpopulation, whereas massive parallel ecDNA reintegration would result in HSRs dispersed across the genome (Figure 1A-B; Supplementary Figure 1A-B).

We first sought to investigate whether ecDNA+ cell populations have reduced transduction efficiency compared to HSR+ cell populations. To directly test the lentiviral transduction efficiency in both cell subpopulations with ecDNA or HSR, we performed single-cell cloning on PC3 cells. We identified two clones consisting of only cells containing extrachromosomal *MYC* amplification (PC3-ecMYC1 and PC3-ecMYC2) and two clones containing chromosomal MYC amplification (PC3-hsrMYC1 and PC3-hsrMYC2) using FISH analysis. We evaluated lentiviral transduction efficiency on these four PC3 single-cell clones using a 9.6kb lentivirus carrying green fluorescent protein (GFP)-expressing plasmid. The single-cell clones were infected with lentivirus for 24 hours and subjected to flow cytometry analysis (**Figure 1F** and **Supplementary Fig. 2A**). We observed that the ecDNA+ PC3 clones have a significantly lower transduction efficiency than those with the same oncogene on HSR (T-test p-value < 0.05). We confirmed the reduced transduction efficiency in ecDNA+ colorectal cells (COLO320DM) after 24-hour lentiviral infection compared to the isogenic pair of the same cells with HSR (COLO320HSR) (T-test, p-value = 0.07) **(Supplementary Fig. 2B)**. Next, we evaluated transduction efficiency in a panel of three ecDNA+ and three HSR+ neuroblastoma cell lines, which consist exclusively of ecDNA+ or HSR+ cells. We observed the same result, with HSR+ neuroblastoma cell lines showing significantly higher transduction efficacy than ecDNA+ neuroblastoma cell lines after 48-hour lentiviral infection (two-way ANOVA, p-value = 0.014) **(Supplementary Fig. 2C)**. These results implicate a lower transduction efficiency as driving a depletion of ecDNA+ cells following lentiviral transduction of a heterogeneous cell population and selection for drug resistance.

Relative depletion of ecDNA+ cells can also be achieved by a difference in the proliferation rate of ecDNA+ cells compared to HSR+ cells. Therefore, we investigated cell growth patterns at different time points of lentiviral treatment in both ecDNA+ (EC) and HSR+ (HSR) PC3 clones (**Figure 1G** and **Supplementary Fig. 3A + B**). The pure ecDNA+ or HSR+ PC3 clones were treated with lentivirus and tested for cell proliferation using CellTiter-Glo assay. The ecDNA+ and HSR+ PC3 clones showed comparable cell proliferation patterns before and after lentiviral transduction, indicating that lentiviral transduction does not alter cell growth patterns.

Next, we analyzed cell death in response to lentiviral transduction. Transduced ecDNA+ cells exhibited higher cell death compared to HSR+ and non-amplified cell lines (Two-way ANOVA, p-value = 0.0493), suggesting that lentiviral transduction can also result in selective lethality in ecDNA+ cells **(Supplementary Fig. 3C)**. Therefore, the selective depletion of ecDNA+ cells following lentiviral transduction appears to result from 1) their lower transduction efficiency, 2) subsequence cell death of non-transduced cells by antibiotics treatment, and 3) the selective lethal effects of the lentivirus.

We assessed the effect of integrase on the depletion of ecDNA+ cells in PC3 and HeLa-MTX-Res cell line models (**Figure 1H**). To achieve this, we used an integrase-defective lentiviral vector (D64V integrase), which mutation showed the strongest inhibition of integrase^9^. Despite the lack of integrase functionality, lentiviral transduction depleted the ecDNA+ populations in both PC3 and HeLa-MTX-Res cell line models, suggesting the absence of a correlation between integrase function and ecDNA depletion.

Finally, we found that transducing PC3 cells using increased infection time (72 hours) resulted in an increase of GFP-expressing cells in ecDNA+ clones compared to HSR+ clones **(Supplemental Figure 4**). This observation suggests that the shorter infection times result in the selection of HSR+ cells, but that this effect can be mitigated by infection times of 72 hours or more. Future experiments are needed to better understand the difference between short-term and long-term infection times.

## Discussion

In this study, we show that lentiviral transduction drives specific depletion of ecDNA+ cells from heterogeneous cell populations. Lentiviral transduction is the most common method for the stable delivery of foreign genes and has, for example, been used in the very popular DepMap resource^10^.Our results suggest that the use of lentivirus requires caution in ecDNA research. To ensure accurate interpretation of such data, thorough validation of ecDNA status will be necessary. Alternative methods to successfully achieve transgene delivery in ecDNA+ cell models are available. First, lentiviral delivery of smaller plasmids did not alter transduction efficiency. Prolonged infection time may result in an increased number of ecDNA+ infected cells. Second, the PiggyBac system did not impact subpopulation fraction shift, indicating that transposon can be used alternately. Transgene delivery using the lentiviral system of a homogeneous ecDNA+ population, i.e., in the absence of HSR+ and other subpopulations, may be able to be successfully achieved by carefully tweaking to overcome differences in transduction efficiency and cell death following transduction.

While our observations suggest a limited use for a common technology in ecDNA research, our results can also be interpreted in a positive way as it may reveal a pathway with the potential to selectively deplete cells driven by ecDNA in cancer.

## Acknowledgments

This work was delivered as part of the eDyNAmiC team supported by the Cancer Grand Challenges partnership funded by Cancer Research UK (CGCATF-2021/100012; CGCATF-2021/100016 to R.G.W.V.; 398299703 to A.G.H.; CGCATF-2021/100018 to J.D.B.) and the National Cancer Institute (OT2CA278688; OT2CA278649 to R.G.W.V.; OT2CA278666 to J.D.B.). This work was also supported by the National Institutes of Health (grants R01 CA237208, R21 NS114873 and R33 CA236681 to R.G.W.V.). A.G.H. is supported by the Deutsche Krebshilfe (German Cancer Aid) Mildred Scheel Professorship program – 70114107. This project has received funding from the European Research Council (ERC) under the European Union’s Horizon 2020 research and innovation programme (grant agreement No. 949172). E.Y. is supported by a grant from the Elsa U. Pardee Foundation.

## Methods

### Cell cultures and cell lines

The human prostate cancer cell line, PC3, was a gift from Dr. Paul Mischel at Stanford University and cultured in F12-K (ATCC, 30-2004) with 10% FBS (VWR, 97068-085). The parental HeLa-S3 cell line was purchased from ATCC and cultured in DMEM (Gibco, 11054020) supplemented with 10% FBS (VWR, 97068-085), 1% penicillin and streptomycin (Gibco, 15140122), and 2mM L-glutamine (Gibco, 25030081). The COLO320DM and COLO320HSR cell lines were purchased from ATCC and cultured in RPMI-1640 (ATCC, 30-2001) supplemented with 10% FBS (VWR, 97068-085). All cultured cells were tested for *mycoplasma* contamination before use with the MycoAlert Mycoplasma Detection Kit (Lonza).

### Generation of HeLa single-cell clones

Parental HeLa S3 cells were treated with 160 nM methotrexate for 2-6 weeks. Resistant cells were harvested and subjected to single-cell cloning. The clones were allowed to expand in low methotrexate concentrations to obtain clonal lines, which were screened by DHFR copy number. Clones with the highest copy number were treated with progressively increasing methotrexate concentrations and characterized by DAPI staining and fluorescence in-situ hybridization analyses of metaphase spreads to identify clones with and without ecDNAs.

### Generation of PC3 single-cell clones

Single PC3 cells were plated in each well of a 96-well plate using a FACSAria cell sorter. Then, each single cell was cultured until a colony was visible. Then, individual colonies from each well were passed into bigger plates and expanded. MYC amplification status of each clone was validated by FISH analysis.

### General lentivirus production and transduction

The 293T cells were seeded onto Poly-L-Lysine (PLL)-coated plates to obtain 70-80% confluence on the following day, on which cells were transfected with packaging and envelope plasmids along with the lentiviral plasmid expressing specific genes using lipofectamine 3000 transfection reagent. The medium was changed 6 hours after transfection. On the next day (29 hours post-transfection), lentiviral supernatants were collected and mixed with Lenti-X Concentrator as per the manufacturer’s instructions. Lentiviral supernatants with concentrator were incubated overnight at 4°C. Lentiviruses were then pelleted using centrifugation, and pellets were re-suspended in PBS; aliquots were prepared and stored at -80°C until use.

For transduction, cells were seeded in 24- or 6-well plates to obtain 60-70% confluence on the following day when they were transduced with indicated lentiviruses using 8 ug/ml polybrene or transfected with Piggybac system plasmids using lipofectamine 3000 transfection reagent. Treatment with a selection marker was started 24 or 48 hours after transfection/transduction. Medium with selection was changed every 2-3 days. Cells were harvested at 1 or 2 weeks after starting selection and processed for metaphase spread preparation and FISH analysis.

### Transduction efficiency test for 72 hour-infection

293T cells were seeded onto cell-culture treated 10cm plates to obtain 70-80% confluence on the following day, on which cells were transfected with packaging and envelope plasmids along with the lentiviral plasmid expressing GFP (pHAGE-EF1α-EGFP-Puro, 9 kb) using lipofectamine 2000 transfection reagent. 48 hours later, lentiviral supernatants were collected, filtered (0.45µM), aliquoted and stored at -80°C until use.

For transduction, PC3 clones were seeded in 6-well plates to obtain 60-70% confluence on the following day (0.1×10^6^ cells/well). The following day cells were transduced with equal amounts of lentivirus. 24 hours post-infection lentiviral media was replenished with fresh media. To assess the infection efficiency, GFP-expressing cells were quantified via flow cytometry 72 hours post-infection.

Cells were washed twice with PBS and resuspended in FACS buffer (PBS with 2% FBS). To ensure a single-cell suspension, samples were passed through cell strainer tubes (Fisher #08-771-23) before analysis. Flow cytometry was performed using a Sony SH800 cell sorter, collecting 100,000 events per sample. GFP fluorescence was detected using the FITC channel (488 nm excitation). Data were analyzed using FlowJo, with GFP-positive cell percentages quantified after gating on live, single-cell populations.

### FISH analysis

PC3 cells with and without lentiviral transduction were treated with 80 ng/mL Colcemid (Roche, 10-295-892-001) for 5 hours (24 hours for HeLa-MTX-Res cells), and harvested to prepare metaphase spreads using standard cytogenetic procedures. Metaphase spreads were subjected to FISH analysis using probes binding to MYC and chromosome 8 (DHFR and chromosome 5 for HeLa-MTX-Res cells). A hybridization buffer (Empire Genomics) mixed with probes (Empire Genomics) was applied to the slides, and the slides were denatured at 75°C for 5 minutes. The slides were then immediately transferred and incubated at 37°C overnight. The post-hybridization wash was with prewarmed 0.4× saline sodium citrate (SSC) at 75°C for 1 minute, followed by a second wash with 2× SSC/0.05% Tween-20 for 2 minutes at room temperature. The slides were then briefly rinsed by water and air-dried. The VECTASHIELD mounting medium with DAPI (Vector Laboratories) was applied and the coverslip was mounted onto a glass slide. Tissue images were scanned under Leica STED 3×/DLS Confocal or Leica Stellaris 5 Confocal with an oil-immersion objective (40×). As excitation laser, 405 nm, 488 nm, and 561 nm were used. Z-stack acquired at a 0.3-to 0.5-μm step size was performed, and all analysis was conducted based on maximum intensity projection images of the 3-D volume of the cells. Images were acquired and processed by LAS X software.

### Real-time qPCR

The total RNA was extracted from ecDNA+ and HSR+ clones using an RNeasy kit (Qiagen) following the manufacturer’s instructions. The RNA quality and concentration were assessed using NanoDrop spectrophotometers (Thermo Scientific). cDNA was synthesized using a High-Capacity cDNA Reverse Transcription kit (Applied Biosystems) according to the manufacturer’s protocol, then a master mix containing SYBR Green PCR Master Mix (Applied Biosystems), forward and reverse primers, and cDNA template was prepared. For the real-time PCR cycling conditions, the initial denaturation was at 95°C for 2 minutes; the denaturation was at 95°C for 15 seconds; the annealing/extension was at 57°C for 30 seconds and at 72°C for 1 minute (repeat for 40 cycles); the melting curve analysis was performed at 95°C for 15 seconds, at 60°C for 1 minute, and at 95°C for 15 seconds. Data analysis was performed using QuantStudio Design & Analysis Software (Applied Biosystems). Ct values were normalized to a reference gene (GAPDH), and relative expression was calculated using the ΔΔCt method.

### Immunofluorescent staining

PC3 clones (1.5 × 10^5^) were plated into a glass-viewing area of a confocal dish (Wuxi NEST Biotechnology, 801002). The next day, cells were briefly rinsed with PBS and fixed with 4% PFA for 10 minutes at RT. The cells were then washed with PBS three times and incubated with the permeabilization buffer (0.1% Triton X-100 in PBS) for 10 minutes, except for the samples for LDLR staining. Then, the cells were incubated with the blocking buffer (1% BSA in 0.1% Tween-20 PBS) for 4 hours at RT. The blocking buffer was then removed, and primary antibodies diluted in the blocking buffer at appropriate concentration were applied (LDLR, ab30532 (1:500); MAP1A, ab101224 (1:100); MAP1S, ab254930 (1:500); TRIM5a, ab190235 (1:500)). The dishes were then incubated with primary antibodies at 4°C overnight. The dishes were then washed with the washing buffer (0.1% Tween-20 in PBS) three times, and the secondary antibody conjugated with Alexa 555 (ab150086, 1:1000) was applied. After an hour of incubation, the dishes were washed three times and counterstained with VECTASHIELD mounting medium with DAPI. The images were scanned under Leica Stellaris 5 Confocal with an oil-immersion objective (40×). As excitation laser, 405 nm, and 561 nm were used. Z-stack acquired at a 0.3-to 0.5-μm step size was performed, and all analysis was conducted based on maximum intensity projection images of the 3-D volume of the cells. Images were acquired and processed by LAS X software.

### Cell viability assay

The cell viability was tested using CellTiter-Glo (Promega) according to the manufacturer’s protocol. The luminescence was measured using the Tecan Infinite 200 Pro microplate reader (Tecan Life Science).

### Flow Cytometry assay

For PC3-derived cells and COLO320 cells, the cells were transduced with lentivirus carrying GFP for 24 hours. For neuroblastoma cell lines, the cells were transduced with lentivirus carrying GFP for 48 hours. The transduced cells were harvested, rinsed with DPBS without calcium and magnesium, and resuspended in FACS buffer (0.1% BSA in DPBS). The number of GFP-positive cells was analyzed using FACSAriaII.

### Statistical analysis

All sample sizes and statistical methods are indicated in the corresponding figure or figure legends. All statistical tests were performed in GraphPad Prism.

## Figure legends

**Supplementary Figure 1.**
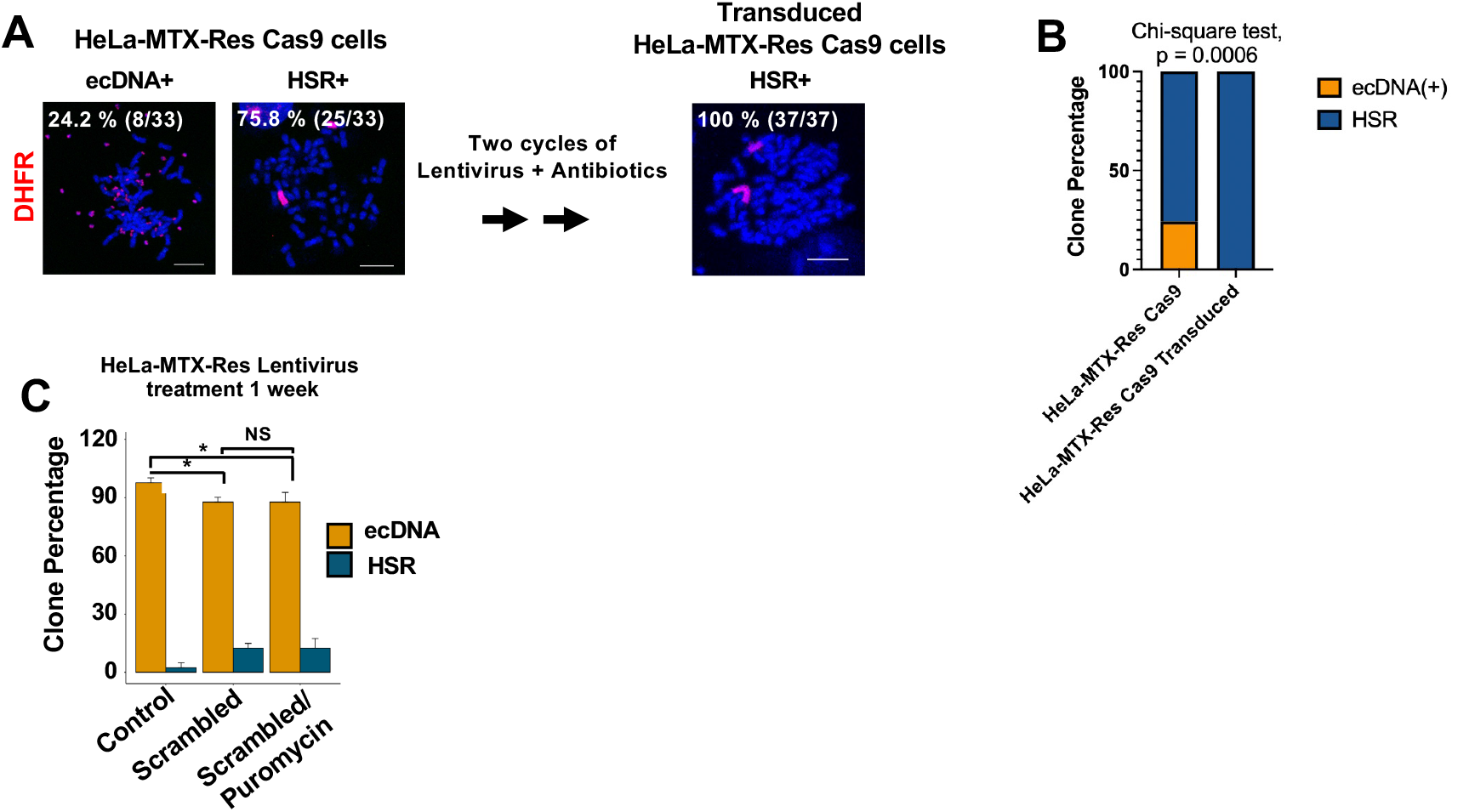
Lentiviral transduction causes the depletion of HeLa cells with extrachromosomally amplified DHFR. **A**. Oncogene amplification status before and after lentiviral transduction. The HeLa cell line that acquired methotrexate resistance (HeLa-MTX-Res) underwent two cycles of lentiviral transduction followed by antibiotics selection. Then, the cells were synchronized at metaphase and processed for FISH analysis. DHFR probe (red) was used. The number of cells containing DHFR amplification was quantified (n = 33). Bar = 10 micrometer. **B**. Graphical summary and statistical test of FISH analysis (Chi-square test, p = 0.0006). **C**. Switch of the subpopulation proportion after a single cycle of lentiviral transduction followed by puromycin treatment. HeLa-MTX-Res cells were lentivirally transduced and selected with puromycin for 1 week. FISH analysis with a DHFR probe was performed to quantify subpopulations (T-test, *p < 0.05, n=3).

**Supplementary Figure 2.**
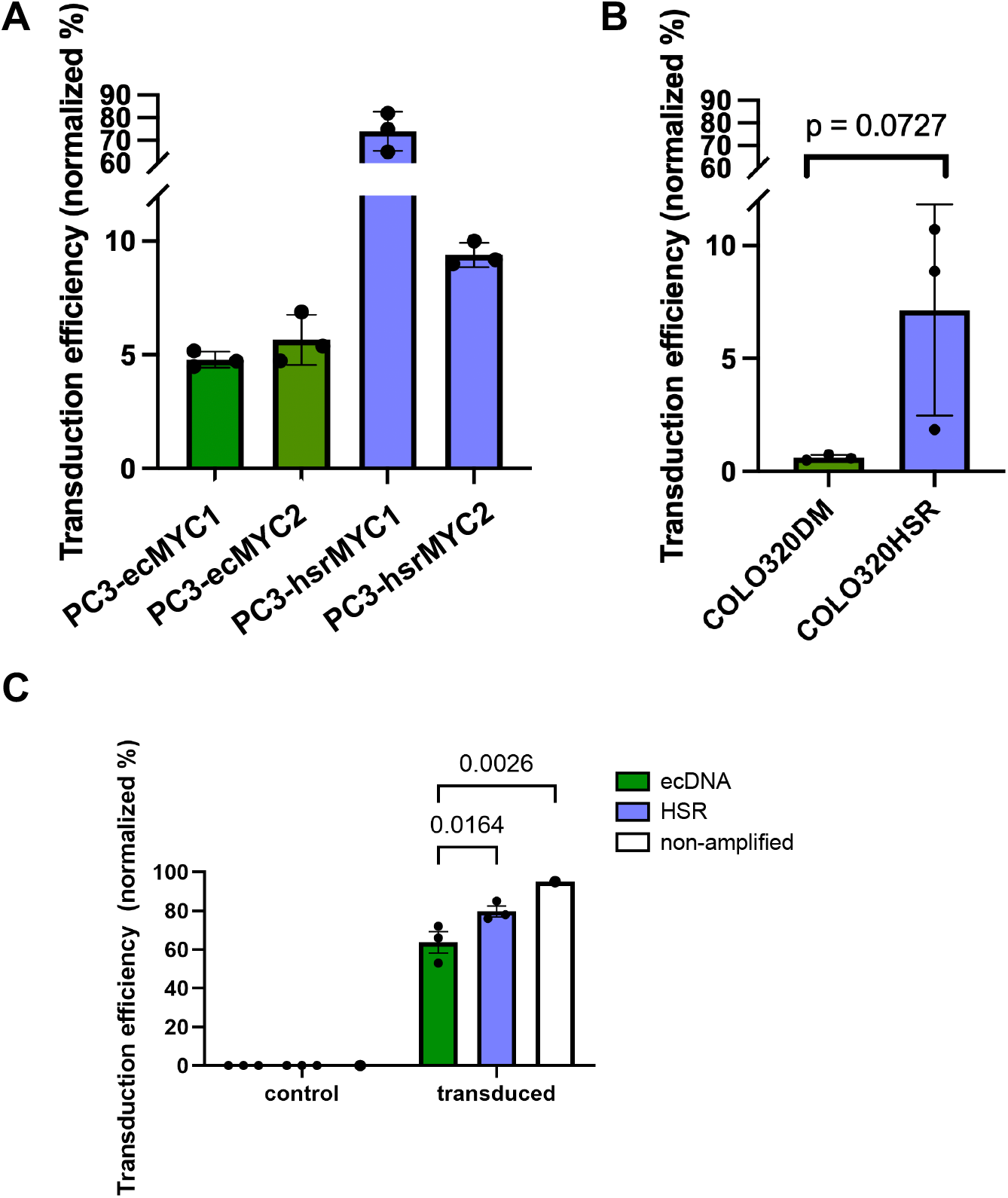
Subpopulations with ecDNA tend to show lower lentiviral transduction efficiency. **A**. Differential lentiviral transduction efficiency in PC3 single-cell clones. Cells were transduced with lentivirus carrying green fluorescence protein (GFP) for 24 hours and subjected to flow cytometry analysis. Lentiviral transduction efficiency was calculated by quantifying the proportion of cells expressing GFP (n = 3, the combined result is shown in Figure 1F). **B**. Differential lentiviral transduction efficiency between COLO320DM (ecDNA+) and COLO320HSR (HSR+). Cells were transduced with lentivirus carrying green fluorescence protein (GFP) for 24 hours and subjected to flow cytometry analysis. Lentiviral transduction efficiency was calculated by quantifying the proportion of cells expressing GFP (T-test, n = 3). **C**. Differential lentiviral transduction efficiency in neuroblastoma cells with ecDNA (CHP-212, TR14, and SiMa) or HSR (IMR-5/75, KELLY, and NGP) *MYCN* amplification or non-amplified (SH-EP). Cells were transduced with lentivirus carrying GFP for 48 hours and analyzed by flow cytometry as above. Significant differences were determined by two-way ANOVA with multiple comparisons.

**Supplementary Figure 3.**
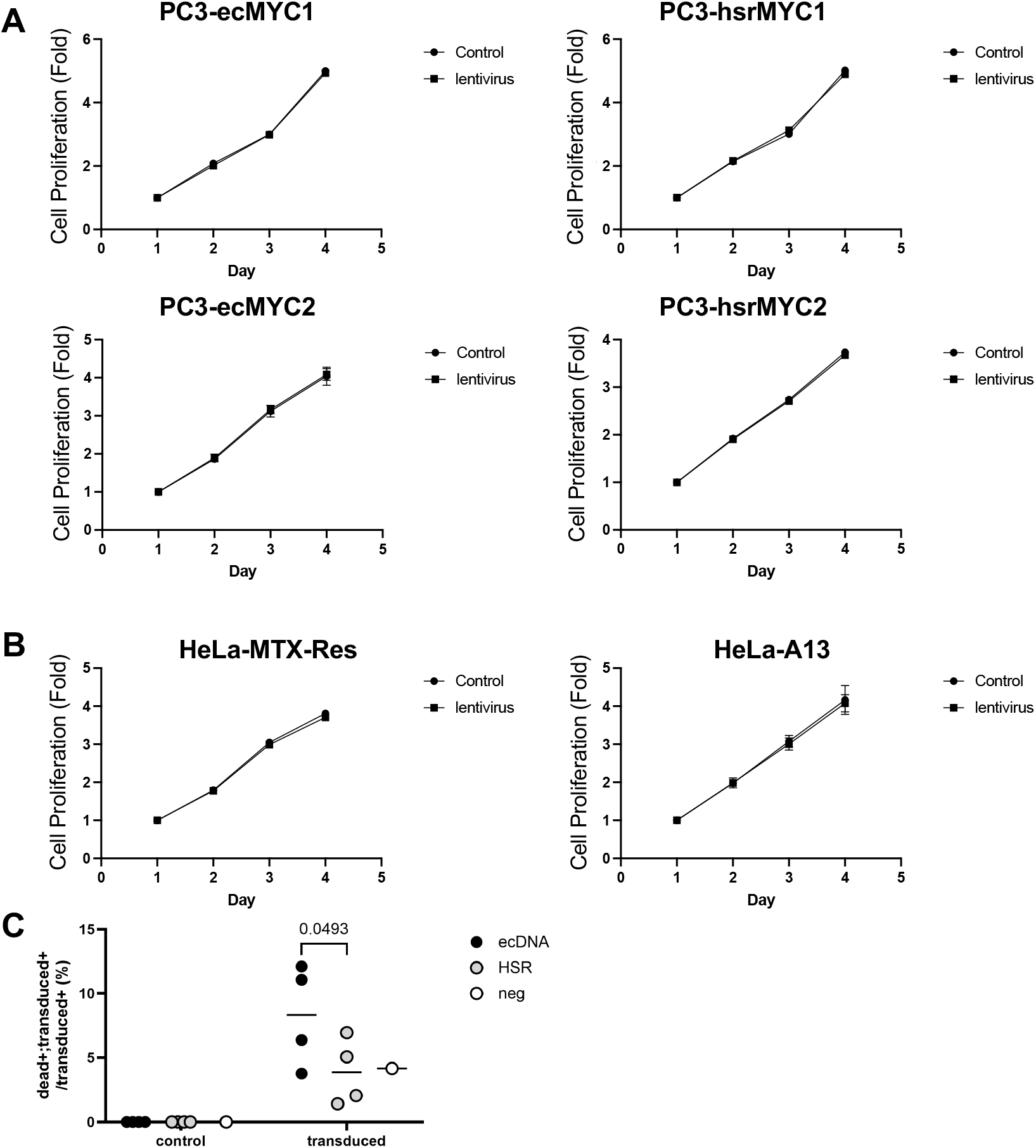
Cell proliferation comparison between cancer single-cell clones with or without ecDNA. **A**. Five sets of each clone were transduced with lentivirus, and each set of cells was subjected to CellTiter-Glo assay on different days (posttransduction 1, 2, 3, 4, 5 days). The posttransduction proliferation rate was compared to the proliferation rate of the untreated control cells (the combined result is shown in Figure 1F). **B**. HeLa-MTX-Res is DHFR-ecDNA+ clones (EC). HeLa-A13 is ecDNA-clones. Five sets of each clone were transduced with lentivirus, and each set of cells was subjected to CellTiter-Glo assay on different days (posttransduction 1, 2, 3, 4, 5 days). The posttransduction proliferation rate was compared to the proliferation rate of the untreated control cells. **C**. Analysis of cell viability in response to lentiviral transduction. ecDNA (CHP-212, TR14, SiMa and COLO320-DM), HSR (IMR-5/75, KELLY, NGP and COLO320-HSR) and non-amplified cells (SH-EP) were transduced with lentivirus carrying GFP and analyzed at 48 hours post-transduction by FACS. Dead cells were identified by 7-AAD staining and transduced cells by GFP signal. Significant differences were determined by two-way ANOVA with multiple comparisons.

**Supplementary Figure 4.**
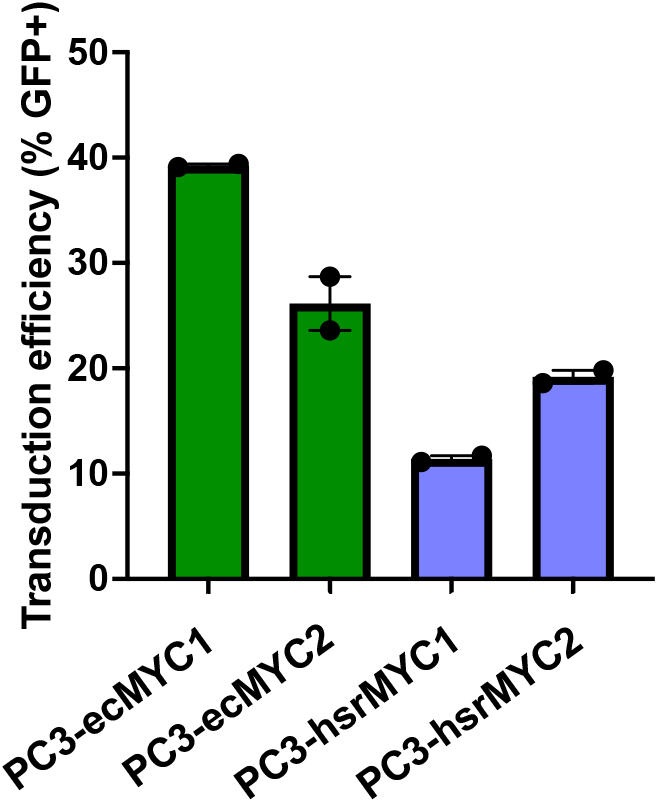
Subpopulations with ecDNA show modestly higher lentiviral transduction efficiency following long-term virus infection. Differential lentiviral transduction efficiency in PC3 single-cell clones. Cells were transduced with lentivirus carrying green fluorescence protein (GFP) for 24 hours and subjected to flow cytometry analysis 72 hours post-infection. Lentiviral transduction efficiency was calculated by quantifying the proportion of cells expressing GFP (n = 2).

